# Transgene by Germplasm Interactions Can Impact Transgene Evaluation

**DOI:** 10.1101/2022.10.22.513364

**Authors:** Julien F Linares, Nathan D Coles, Hua Mo, Jeff E Habben, Sabrina Humbert, Carlos Messina, Tom Tang, Mark Cooper, Carla Gho, Ricardo Carrasco, Javier Carter, Jillian Wicher Flounders, E Charles Brummer

## Abstract

Transgenes have been successfully commercialized for qualitatively inherited insect and herbicide resistance traits that show similar effects across genetic backgrounds. However, for quantitative traits like yield, genetic background may affect the measured transgene value. In this paper, we evaluated whether different genetic backgrounds impact the estimated value of a transgene for grain yield, ear height, and anthesis-silking interval for maize by developing isogenic pairs of lines with and without a transgene and testing them in hybrid combination with non-transgenic lines from a complementary heterotic group across eleven environments in the USA. Over all hybrid combinations, the transgene increased yield by 0.2 Mg ha^−1^. Across multiple non-transgenic lines of the opposing heterotic group, the transgene effect within a line pair ranged from an increase of 0.8 Mg ha^−1^ for the NSS4 and SS7 transgenic lines to a reduction of 0.3 Mg ha^−1^ for the NSS5 transgenic line when compared to their non-transgenic isoline. Transgenic hybrids were often taller than non-transgenic hybrids (P<0.05). Anthesis to silking interval was reduced by 4□C growing degree units overall, but no transgene × genotype interaction was detected among line pairs. Our results show the importance of testing transgene efficacy across a large sample of elite hybrid pairs to assess the gene’s value. By only testing in a specific hybrid background, as may be done for qualitative traits like insect resistance, transgenes could be erroneously advanced or eliminated.

## INTRODUCTION

Plant breeding programs have been highly successful at improving yield and other characteristics of maize and other crops (Duvick et al. 2004, Smith et al. 2014, Andorf et al., 2019). Since the mid-1990s, commercial maize hybrids containing transgenes to help protect against pests or be resistant to herbicides have been developed. These hybrids targeted a specific gene to directly produce a large effect on the desired trait phenotype. This approach has had tremendous commercial success, aiding farmers to deliver ever increasing yields on a yearly basis while maintaining profitability and minimizing environmental impact (Brookes and Barfoot, 2017). However, using transgenes to improve complex traits, such as yield, has proven to be a challenging task (Guo et al. 2014, Simmons et al. 2021). For yield enhancement, the complexity of gene interactions across the genome has made successful transgene discovery and evaluation extremely difficult. Many studies have claimed to identify quantitative trait loci (QTL) or transgenes that promised to significantly increase yield (Nelson et al. 2007; Chen et al., 2017), yet most single yield transgenes have not materialized into successful commercial products (Nuccio et al., 2018, Simmons et al. 2021). Part of the reason behind this phenomenon is the complex genetic architecture underlying highly quantitative traits (gene × gene, genotype × transgene, and genotype × transgene × environment interactions). Although these interactions can also present challenges for pest and herbicide resistance transgenes (Elmore et al., 2001), the primary target of these transgenes produces a dominant and highly penetrant phenotype across germplasm. While these pest and herbicide transgenes have been selected to have direct effects on their intended targets, they can also have pleiotropic effects on complex traits like yield in the absence of the primary stressor (Xia et al., 2010; Darmency, 2013).

Testing the effect of a single transgene or QTL on a quantitatively inherited trait phenotype is difficult when comparing lines from different genetic backgrounds. To minimize the impact of the genetic background, near-isogenic lines (NILs) are typically used for comparison. But because NILs typically require three or more generations of backcrossing to recover at minimum 95 percent for the recurrent parent (Frisch, 1999), the development of a diverse set of germplasm in which to evaluate a gene of interest is typically not time or cost effective, and hence, not done (e.g., Simmons et al. 2021). Forward breeding, whereby breeding populations in which the transgene has been integrated are developed and selected, could overcome some of these challenges. However, given the current complex regulatory landscape, the time taken, and the cost of developing populations with non-approved transgenic events, forward breeding has not been considered an appealing approach (Mumm, 2007).

Therefore, a challenge facing breeders is how to best evaluate transgenes to accurately quantify their average impact on complex traits across a diverse range of elite breeding populations. Without an experimental system to generate these data, optimized selection of genetic backgrounds that minimize yield drag or that maximize yield enhancement cannot be achieved. Consequently, some transgenes that might have significant benefits may never be pursued for commercialization because the narrow germplasm in which they were tested did not show their potential advantages.

In this study, we consider a transgene that influences yield through its role in ethylene biosynthesis to investigate transgene × genotype × environment interactions for yield and other important agronomic traits. Ethylene, a volatile hormone, has a significant and diverse impact on plant growth and development, from the induction of reproduction to biotic and abiotic stress responses (Iqbal et al., 2017; Taiz and Zeiger, 2002). With such a broad range of impact, altering ethylene synthesis can cause multiple non-targeted, secondary, downstream effects. The ethylene biosynthetic pathway is rather simple, and reduction of ethylene evolution has been achieved in multiple ways from chemical applications to gene modifications (Shi et al., 2016; Khan et al., 2008; Jackson et al., 1981). Ethylene is synthesized in plants from methionine. The first step of the process consists of methionine being converted to S-adenosyl-L-methionine (SAM) by SAM synthase. SAM is then converted to 1-aminocyclopropane-1-carboxylic acid (ACC) by ACC synthase (ACS), and finally, ethylene is synthesized from ACC by ACC oxidase (Yang and Hoffman, 1984). The major rate limiting step to this pathway is the conversion of SAM to ACC by ACS. Manipulation of this rate limiting step through the introduction of an ACS6 RNAi transgene reduced ethylene evolution by an average of 50% across many different transgene integration events and, in a limited number of maize hybrids, it increased grain yield under water limiting conditions (Habben et al., 2014).

In this experiment, we evaluated a single event of the ACS6 RNAi transgene across multiple hybrids and their respective NILs. We hypothesized that the effect of the transgene on grain yield and other agronomic traits would vary across genetic backgrounds. The objective of this experiment was to test this hypothesis in order to quantify the importance of testing diverse hybrids when determining transgene effects on agronomically important traits in elite maize breeding programs.

## MATERIALS AND METHODS

### Germplasm

Eleven elite non-transgenic maize inbred lines from the commercial Corteva germplasm pool and their respective near-isogenic transgenic lines all containing the same ACS6 RNAi event were used to generate hybrids for experimental testing. For ten of these inbred lines, backcrossing, using foreground and background selection with genetic markers, was used to introgress the event with at least three backcross generations, resulting in near-isogenic lines with at least 95 percent of the recurrent parent. After the final backcross, lines were self-pollinated and individuals homozygous for the transgene event were selected. The 11^th^ transgenic inbred line (SS6) was the result of direct transformation. Of the 11 inbred line pairs, five were from the Stiff Stalk (SS) heterotic group and six from the Non-Stiff Stalk (NSS) heterotic group.

A total of 124 hybrids were created for inclusion in the experiment. Each pair of lines (non-transgenic and transgenic) from one heterotic group was crossed to the non-transgenic lines from the complementary heterotic group. This crossing scheme generated a ‘nest’ of three hybrids for any two inbred lines crossed (Table 1). Of these three hybrids, two were hemizygous, with the transgene being introduced by the transgenic inbred from either the SS or NSS heterotic group; and one was non-transgenic. The homozygous transgene hybrid was not created. Therefore, by crossing the five inbred line pairs (10 inbred lines total) from the SS heterotic group with the six pairs (12 inbred lines) from the complementary NSS heterotic group, we generated 90 hybrids. To expand the germplasm tested, an additional two SS and one NSS non-transgenic inbred lines were crossed to all pairs of lines (non-transgenic and transgenic) from the opposite heterotic group, generating an additional 34 entries. In all 124 hybrid combinations, the SS line was used as the female parent.

**Table 1.**
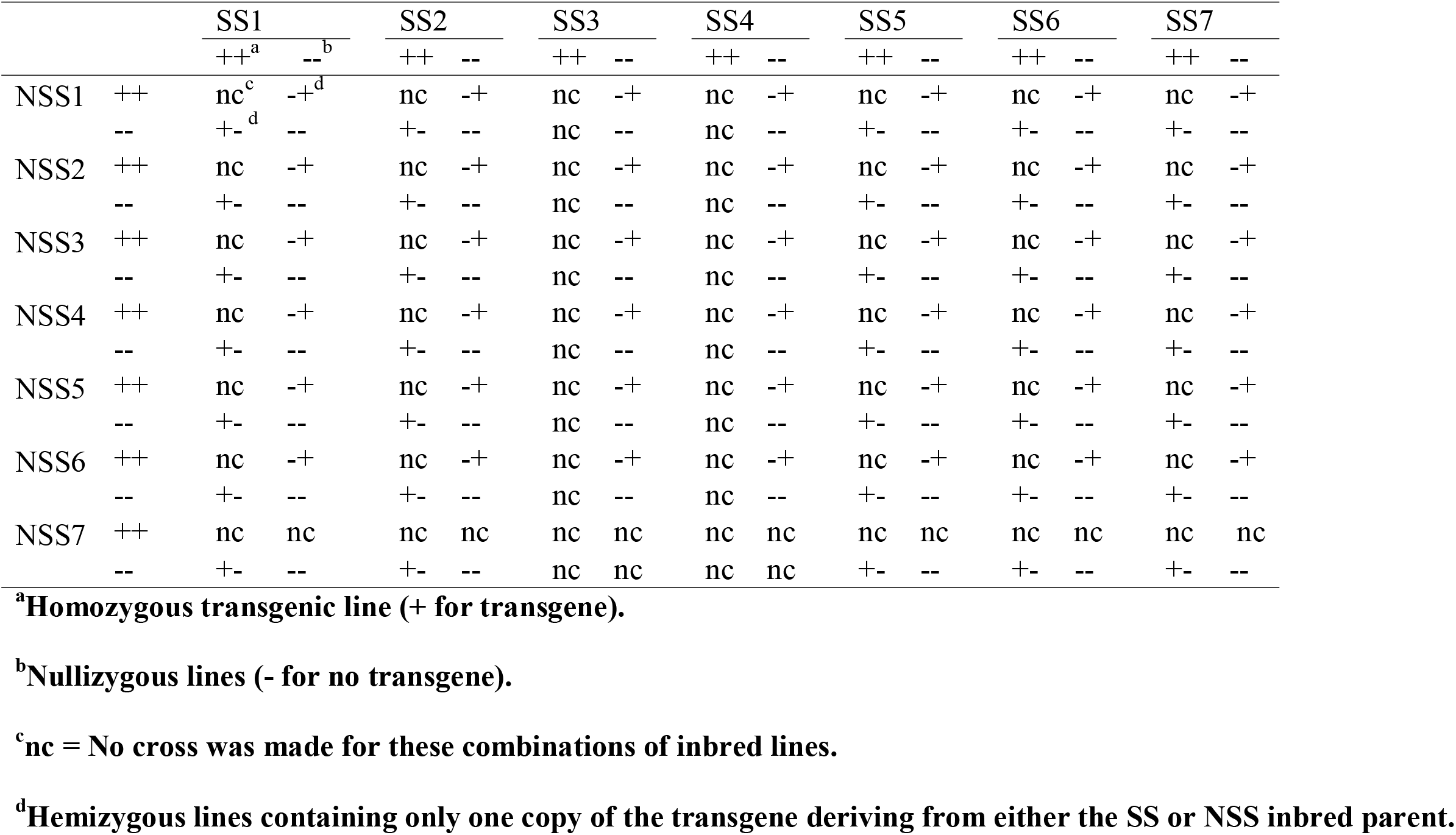
Crossing scheme used to generate hybrids evaluated in this experiment. Hybrid combinations were generated by crossing stiff stalk (SS) inbred lines to non-stiff stalk inbred lines (NSS). Zygosity is depicted as “+-” for hemizygous hybrids where the transgene donor was a SS, “-+” for hemizygous hybrids where the transgene donor was a NSS, and “--” for nullizygous hybrids. “nc” indicates no hybrid seed was produced for that cross.

### Environments evaluated

Field trials were conducted at eleven locations across the United States in 2017 and 2018 (Table 2). These locations included both fully irrigated and managed water-stress environments. Other than the imposed water limitations, all environments followed standard agronomic practices for soil preparation and plant growth. All experimental plots consisted of 2-rows separated by 0.75 m and were 2.4 m long. Border plots were planted around the outer edges of the experiment to reduce edge effects. The target plant population for all plots was 95,000 plants per hectare. The experimental design used at all locations was a split-plot, where the main-plot treatment was the hybrid nest (e.g., the set of three hybrids derived from SS1 × NSS1, or a set of two hybrids if one hybrid parent did not have a transgenic version) and the split-plot treatment included the three (or two hybrids) versions of that hybrid (i.e., +/−, −/+, and −/−) with two to four replications per location. Managed water-stress trials were conducted at Woodland, CA and Plainview, TX, which both had little to no rainfall during the growing season. Water-stress for those locations was targeted at the flowering stage by withholding irrigation during the flowering period of the hybrids within the experiments. The managed water-stressed environments had yield reductions of 40-60% and an average increase in anthesis to silking interval (ASI) of 19.4 Celsius growing degree units (GDUs) compared to well-watered environments.

**Table 2.** All locations utilized in experimental testing. Soil type as indicated by the USDA Natural Resource Conservation Service. The last letter in the location name indicates whether a locations was considered water-stressed (‘S’) or well-watered (‘W’) in the environmental clusters.

| Location Name | City, State | Planting Date | Harvest Date | Soil Type |
| --- | --- | --- | --- | --- |
| WLD17IW | Woodland, CA | 05/03/2017 | 09/09/2017 | Brentwood Silty Clay Loam |
| WLD17JW | Woodland, CA | 05/02/2017 | 09/02/2017 | Yolo Silt Loam |
| WLD17JS | Woodland, CA | 05/24/2017 | 10/02/3017 | Yolo Silt Loam |
| WLD177S | Woodland, CA | 05/20/2017 | 09/18/2017 | Yolo Silt Loam |
| WLD177W | Woodland, CA | 05/20/2017 | 09/20/2017 | Yolo Silt Loam |
| JST184W | Johnston, IA | 04/27/2018 | 09/17/2018 | Wiota Silty Clay Loam |
| MRN182W | Linn, IA | 05/07/2018 | 10/03/2018 | Franklin Silt Loam |
| WLD189W | Woodland, CA | 04/30/2018 | 09/15/2018 | Yolo Silt Loam |
| WLD189S | Woodland, CA | 04/30/2018 | 09/06/2018 | Yolo Silt Loam |
| WLD18S | Esparto, CA | 05/18/2018 | 09/20/2018 | Brentwood Silty Clay Loam |
| PVW181S | Hale, TX | 04/30/2018 | 09/02/2018 | Pullman Clay Loam |

### Phenotypes collected

Plant height was estimated from the average of four plants subsampled within each plot at R3 and was measured from the ground surface to the node of the flag leaf. Days to anthesis and silking were estimated based on the date when 50% of the plants in the plot had any anthers shedding pollen or silks extruded from the sheath, respectively. Days to silking and anthesis were then converted to growing degree units (GDUs) for the analysis. Anthesis to silking interval was calculated by taking the difference between GDUs to silking and GDUs to anthesis. All grain yield measurements were obtained with a standard two-plot combine. For each plot all plants were harvested, the total kernel weight of the plot was obtained, and grain moisture was determined. The plot weight was then converted to grain yield as Mg ha^−1^ at 15% moisture.

### Statistical analysis

For the purposes of analysis, the hybrid nests were treated as a design factor and considered to be a “germplasm group.” The analysis considered the transgene effects both overall across hybrids and separately for each heterotic group.

Grain yield, ear height, and anthesis-silking interval were analyzed using mixed linear models. One model was used to predict effects of the transgene and another model was used to examine transgene × germplasm effects, both following the same general form of the following linear mixed model (with specific models in ASREML format shown in Appendix 1):

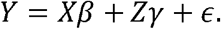

where *Y* is the response variable, and the matrices 1. and’S are the design matrices for fixed and random effects, respectively. We assume *E*(*ϵ*|*X*, *Z*) = 0 and *Cov*(*ϵ*|*X*, *Z*) = *R* with heterogenous autoregression spatial structure (Gilmour et al. 1997, Cadena et al., 2000). *R* is a matrix with the elements in row *i* and column *j* Satisfying *γ_ij_* = *σ_i_σ_j_ρ*^|*i*−*j*|^ 0 < *ρ* < 1. For the random effects, we also assumed that *E*(*γ*) = 0, *Cov*(*γ*, *ϵ*) = 0 and the variance of the parameter vector *γ* was estimated using ASReml. All tests (including main effects and random effects) are performed at the significance level of 5%. The vector *β* represents the fixed effects including transgene, transgenic donor at the heterotic group level (SS or NSS), and NSS and SS parents. Unbiased estimates 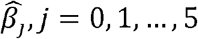 the fixed effects were calculated. The transgene’s effect among transgenic lines across non-transgenic lines, and among non-transgenic lines across transgenic lines were estimated using contrasts between appropriate transgenic and non-transgenic SS and NSS parents. The model included 13 random effects, where *Z* = [*Z*_1_, *Z*_2_, *Z*_3_,…] is used to represent their design matrix, while the corresponding parameter *γ* = (*γ*’_1_, *γ*’_2_, *γ*’_3_,…)’ is predicted using best linear unbiased predictions (BLUPs). The random effects include environments; germplasm; replication within an environment; the transgene × environment, transgene × environment × heterotic group, transgene × germplasm, transgene × germplasm × heterotic group, environment × germplasm × replication, environment × germplasm, transgene × germplasm × environment, and transgene × germplasm × environment × heterotic group interaction effects; and within environment column and row effects. To model transgene × germplasm effects, we dropped the NSS and SS parent line effects to capture all the variance in the transgene × germplasm term. The details for the models can be found in Appendix A. A Z-ratio was calculated by dividing each estimated 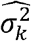 by its own estimated standard error in order to compute the significance of each variance component.

We declared transgene × germplasm and transgene × germplasm × location interactions significant if the z-ratio for the random interaction effects was greater than 1.96.

We evaluated results overall for the transgene, for the transgene by heterotic group (NSS and SS), among transgenic lines across non-transgenic lines, among non-transgenic lines across transgenic lines and by hybrid nest. To predict the effects of the transgenic lines we averaged across all non-transgenic lines (e.g., the average effect of the line pairs of the NSS1 transgenic line across the SS1, SS2, SS3, SS4, SS5, SS6, SS7 non-transgenic lines) while for predicting the impact of non-transgenic lines across transgenic lines we averaged across transgenic lines (e.g. the average effect of line pairs of the non-transgenic NSS1 across the SS1, SS2, SS5, SS6, SS7 transgenic lines).

Prior evidence investigating the efficacy of the ACS6 RNAi transgene indicated it was expected to improve yield in water-stressed environments without decreasing yield under well-watered conditions compared to existing elite hybrids. Hence, we classified the environments as either water-stressed (5 environments) or well-watered (6 environments) prior to running our analyses (Table 2). We initially analyzed all environments together and subsequently conducted separate analyses of yield and plant height for water-stressed and well-watered environments due to the presence of T × G × E interactions.

Mean separations were made using 95% confidence intervals. All statistical significance was assessed at the 5% level. Statistical analyses was done using ASREML (Cadena et al., 2000).

## RESULTS

### Across all genetic backgrounds

Across all environments and germplasm, hybrids containing the transgene yielded 0.2 Mg ha^−1^ more, measured 4 cm taller, and had an estimated ASI of 4 □C GDU less than hybrids without the transgene (Table 3). Results for yield and plant height varied by location and genetic background while the transgene’s effect on ASI was penetrant with no interaction with either location or genetic background. In well-watered environments, the transgene had no significant effect on yield but continued to increase plant height by 5 cm. In the stressed environments, the transgenic hybrids yielded 0.2 Mg ha^−1^ more and measured 3 cm taller than the non-transgenic hybrids.

**Table 3.** Best Linear Unbiased Predictors (BLUP) of hybrid performance for non-transgenic, non-stiff stalk (NSS) transgenic and stiff-stalk (SS) transgenic inbred maize lines when crossed to a non-transgenic line of the opposite heterotic group for grain yield, plant height and ASI traits across all locations, flowering stress (FS) locations only, and well-watered (WW) locations only.

| Lines | Yield |  |  | Plant Height |  |  | ASI<sup>a</sup> |  |  |
| --- | --- | --- | --- | --- | --- | --- | --- | --- | --- |
|  | Overall | FS<sup>b</sup> | WW<sup>c</sup> | Overall | FS | WW | Overall | FS | WW |
|  | -----Mg ha<sup>-1</sup>----- |  |  | -----cm----- |  |  | -----°C----- |  |  |
| Non-Transgenic | 10.0 a<sup>d</sup> | 7.8 a | 11.9 a | 266 a | 247 a | 284 a | 11 a | 24 a | -1 a |
| Trangenic NSS<sup>e</sup> | 10.2 b | 8.0 ab | 12.1 a | 270 b | 250 b | 290 c | 8 b | 19 b | -4 a |
| Transgenic SS<sup>f</sup> | 10.3 b | 8.2 b | 12.0 a | 269 b | 250 b | 288 b | 7 b | 19 b | -5 a |

<sup>a</sup>ASI = anthesis to silking interval, <sup>b</sup>FS=Flowering stress, <sup>c</sup>WW=Well-watered, <sup>d</sup>Means within columns followed by different letters are statistically significant based on Fisher's protected LSD at the 5% probability level, <sup>e</sup>NSS=Non Stiff Stalk, <sup>f</sup>SS=Stiff Stalk.

### By heterotic group donors

Hybrids created from the NSS transgenic lines yielded 0.2 Mg ha^−1^ more than non-transgenic hybrids, while the hybrids created from the SS transgenic lines increased yield by 0.3 Mg ha^−1^ (Table 3) across all environments. Within stress environments only the transgenic hybrids created from the SS transgenic lines increased yield, and this was by an average of 0.4 Mg ha^−1^. Within the well-watered environments there were no significant effects for either SS or NSS transgenic hybrids.

Across all environments, NSS transgenic hybrids were 4 cm taller, and SS transgenic hybrids were 3 cm taller than non-transgenic hybrids (Table 3). In both environments, the transgenic hybrids were taller than the non-transgenic hybrids. In stressed environments, the transgenic hybrids of both heterotic groups increased plant height compared to non-transgenic hybrids, but the transgene effect did not differ among heterotic groups. Under well-watered environments plant height was taller in both NSS and SS transgenic hybrids in comparison to non-transgenic hybrids; NSS and SS transgenic hybrids also differed from one another with the SS transgenic hybrids measuring 4 cm taller while the NSS transgenic hybrids measured 6 cm taller than the non-transgenic hybrids (Table 3).

### By line pair results

Although T x E was not significant for any trait, the T x G and T x G x E interactions were significant for both plant height and yield; hence, the transgene effect was estimated independently for each line pair by comparing the testcross hybrids for a given transgenic isogenic line with the respective hybrids from its corresponding non-transgenic isogenic line. For example, the transgenic hemizygous hybrids developed by crossing transgenic NSS1 to all the SS non-transgenic lines were compared to hybrids from the non-transgenic NSS1 crossed to the same set of SS lines. ASI showed no interactions and hence no additional results are presented below for this trait.

Across all environments and averaged across non-transgenic lines of the opposite heterotic group, the transgenic isogenic line increased yield over the non-transgenic isogenic line for three of eleven line pairs (by 0.8 Mg ha^−1^ for NSS4 and SS7, and 0.3 Mg ha^−1^ for NSS2), and reduced yield by 0.3 Mg ha^−1^ for NSS5 (Table 4). The transgenic and non-transgenic isolines were not different for seven of the line pairs (Table 4). In the stress environments, the transgene decreased yield for the NSS5 line pair, increased yield for the NSS2, NSS4, SS6, and SS7 line pairs, and did not affect yield for the other lines. No yield differences were observed under non-stress environments. Comparison of the transgene effects among NSS line pair hybrids showed a range of 1.1 Mg ha^−1^ for yield across all environments, of 1.6 Mg ha^−1^ in stress environments, and no differences in well-watered environments. Among the hybrids of the SS lines, the range of effects of the transgene was 0.9 Mg ha^−1^ overall, 0.7 Mg ha^−1^ in water stressed environments, and not present in the well-watered environment grouping (Table 4).

**Table 4.** The effect of the transgene (positive numbers indicate the transgene increased the trait value) on yield and plant height for each set of line pairs across all non-transgenic lines of the opposite heterotic group over all environments and within flowering stress (FS) and well-watered (WW) environments.

| Line Pair | Yield |  |  |  |  |  | Plant Height |  |  |  |  |  |
| --- | --- | --- | --- | --- | --- | --- | --- | --- | --- | --- | --- | --- |
|  | Overall |  | FS<sup>a</sup> |  | WW<sup>b</sup> |  | Overall |  | FS |  | WW |  |
|  |  |  | -----Mg ha<sup>-1</sup>----- |  |  |  |  |  | -----cm----- |  |  |  |
| NSS1<sup>c</sup> | 0.1<sup>NS d</sup> | ab<sup>e</sup> | 0.0<sup>NS</sup> | ab | 0.2<sup>NS</sup> | a | 5 * | b | 4 * | b | 6 * | b |
| NSS2 | 0.3 * | b | 0.4 * | b | 0.2<sup>NS</sup> | a | 6 * | b | 4 * | b | 9 * | b |
| NSS3 | 0.3<sup>NS</sup> | b | 0.2<sup>NS</sup> | b | 0.4<sup>NS</sup> | a | 6 * | b | 4 * | b | 8 * | b |
| NSS4 | 0.8 * | c | 1.1 * | c | 0.3<sup>NS</sup> | a | 4 * | b | 3 * | b | 5 * | b |
| NSS5 | -0.3 * | a | -0.5 * | a | -0.1<sup>NS</sup> | a | -3 * | a | -2 * | a | -3<sup>NS</sup> | a |
| NSS6 | 0.1<sup>NS</sup> | ab | 0.0<sup>NS</sup> | ab | 0.4<sup>NS</sup> | a | 6 * | b | 4 * | b | 9 * | b |
| SS1 | -0.1<sup>NS</sup> | a | 0.1<sup>NS</sup> | a | -0.2<sup>NS</sup> | a | 0<sup>NS</sup> | a | 1<sup>NS</sup> | a | 0<sup>NS</sup> | a |
| SS2 | 0.1<sup>NS</sup> | a | 0.2<sup>NS</sup> | a | -0.1<sup>NS</sup> | a | 3 * | bc | 3 * | b | 3<sup>NS</sup> | a |
| SS5 | 0.3<sup>NS</sup> | a | 0.4<sup>NS</sup> | ab | 0.2<sup>NS</sup> | a | 5 * | c | 4 * | b | 8 * | b |
| SS6 | 0.2<sup>NS</sup> | a | 0.5 * | ab | 0.2<sup>NS</sup> | a | 2 * | ab | 0<sup>NS</sup> | a | 5 * | ab |
| SS7 | 0.8 * | b | 0.8 * | b | 0.7 * | a | 3 * | bc | 4 * | b | 2<sup>NS</sup> | a |

<sup>a</sup>FS=Flowering stress, <sup>b</sup>WW=Well-watered, <sup>c</sup>NSS=Non Stiff Stalk, SS=Stiff Stalk, <sup>d</sup>Difference between transgenic and non-transgenic hybrids within a given line pair is non-significant (NS) or significant (\*) at the 5% probability level, <sup>e</sup>Means within columns for each heterotic group (NSS or SS) followed by different letters are statistically significant based on Fisher's protected LSD at the 5% probability level.

We also evaluated the average transgene effect observed when a given non-transgenic line was crossed to line pairs of the other heterotic group. Across all environments, five NSS and three SS non-transgenic lines showed transgene effects across line pair hybrids (Table 5). As for the lines pairs analysis (Table 4), large differences in the effect of the transgene was seen for both SS and NSS non-transgenic lines (Table 5). The NSS non-transgenic lines showed a transgene effect when crossed to SS line pairs ranging from 0.7 Mg ha^−1^ for NSS4, which was 0.6 Mg ha^−1^ more than the effect seen for NSS5. The transgene effect for the SS7 non-transgenic line was greater than for SS1, SS2, SS5, or SS6. The two SS lines that did not have transgenic versions both showed a positive transgene effect when crossed to NSS line pairs. In stress environments, NSS1, NSS3, NSS4, NSS6, NSS7 non-transgenic lines once again showed a transgene effect when crossed to SS line pairs. The transgene effect for the NSS4 non-transgenic line was larger than all other NSS non-transgenic lines, at 1.0 Mg ha^−1^. Among SS non-transgenic lines, only SS4, which was only included as a non-transgenic line, showed a transgene effect when crossed to NSS line pairs, with a difference of 0.5 Mg ha^−1^ between transgenic and non-transgenic hybrids. In the well-watered environments, no differences were detected for NSS or SS non-transgenic lines across transgenic line pairs.

**Table 5.**
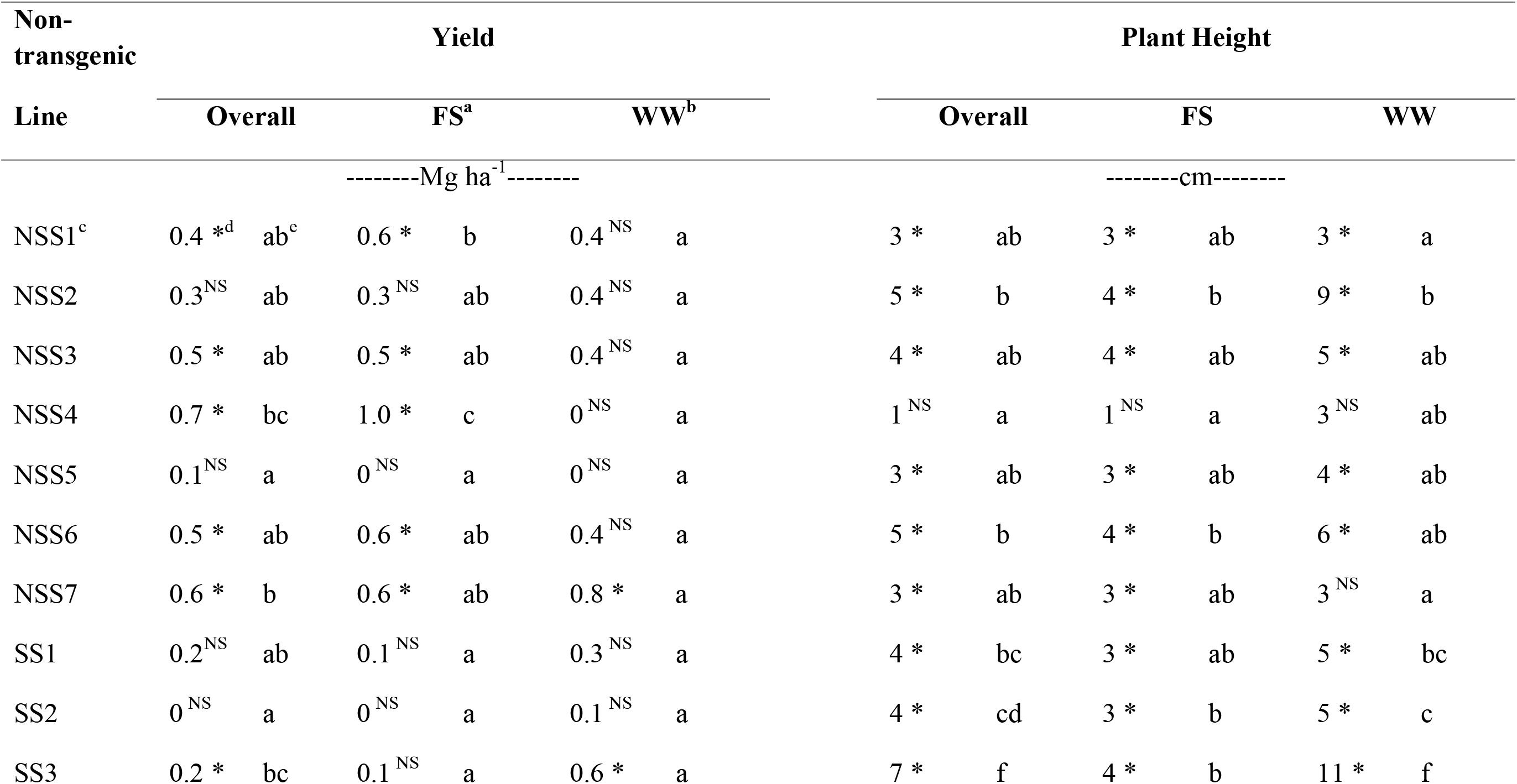
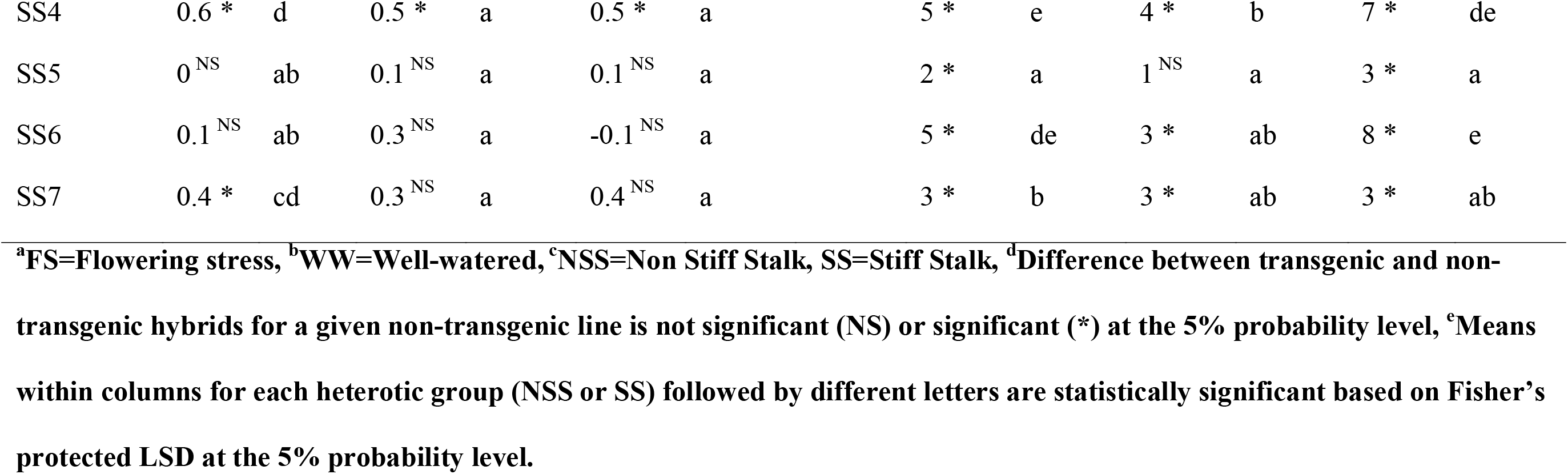
The effect of the transgene (positive numbers indicate the transgene increased the trait value) on yield and plant height of hybrids of non-transgenic (NT) lines across line pairs of the opposite heterotic group over all environments and within flowering stress (FS) and well-watered (WW) environments.

In several cases, both line pairs crossed to non-transgenic lines and the non-transgenic isoline crossed to line pairs showed similar transgene effects. For example, NSS4, when crossed to SS non-transgenic lines, showed a large line pair effect for yield (Table 4) and the non-transgenic NSS4 showed a similarly large effect when crossed to SS line pairs (Table 5). A similar pattern was observed for SS7, except that the non-transgenic line did not show an effect under stress (Table 5). In some cases, a line pair showed no effect (e.g., NSS6) but the non-transgenic isoline of the pair showed a large transgene effect when crossed to SS line pairs. In general, more differences were noted for non-transgenic lines than for line pairs.

The transgene’s effect on plant height was more penetrant than for yield, yet continued to show variation among line pairs across non-transgenic lines (Table 4) and among non-transgenic lines across line pairs (Table 5). Across all environments, the transgenic line of a line pair produced taller hybrids than the non-transgenic line, when crossed to a non-transgenic line of the opposite heterotic group (Table 4), except for SS1 (no difference) and NSS5 (3 cm shorter). Among NSS line pairs, only NSS5 differed from the others, with a negative transgene effect. In both water stressed and fully irrigated environments, line pairs showed a similar effect, with a positive transgene effect on height, with several exceptions (Table 4).

### By hybrid results

Most individual combinations of specific SS and NSS lines did not show differences in yield between transgenic and non-transgenic hybrid performance across all locations, in well-watered environments, or in stressed environments (Supp. Table 1). This is likely due to the small number of data points for each combination. Spearman rank correlations of hybrid yield performance between the two environments was zero. More differences were noted for plant height, and the rank correlation for hybrid height between the two environments was 0.5.

## DISCUSSION

The evaluation of transgene effects is typically done on only one or a small subset of lines initially. This approach has been widely successful for qualitative genes targeting pests and herbicide tolerance with over 400 events submitted in maize alone for deregulation sowed in over 190 million hectares worldwide (ISAAA, 2018). In contrast, fewer than 10 events have been submitted for deregulation for abiotic stress tolerance, yield, or growth genes with a planted acreage of less than 1% of the total area (ISAAA, 2018). Through a review of Corteva’s biotech program intended to impact agronomic traits, including yield, transgene by germplasm interactions were often cited as significant (Simmons et al., 2021). A possible reason for the lack of quantitative trait transgenes is the risk of confounding the estimated effects with genetic background whenever transgene by genetic background effects are present. To assess the potential influence of transgene x genetic background effects for an event of an AC6 RNAi transgene, we evaluated several traits in multiple hybrids across multiple environments. Interactions among transgene and germplasm (T x G) and transgene, germplasm, and environment (T x G x E) were present for plant height in both stress and non-stress environments and for yield across all environments. The transgene’s constitutive effect, reducing ethylene production across all stages of plant development, effected yield and agronomic traits.

The effect of the transgene on yield varied across environments and hybrid background. When analyzing line pairs among non-transgenic lines across transgenic lines, or when evaluating line pairs among transgenic lines across non-transgenic lines, substantial differences in the transgene effect were noted for yield and for plant height. Large rank changes in transgene effects by hybrid were observed between water-stressed and well-watered environments for yield and plant height. Many individual hybrids were not significantly different from their non-transgenic comparators. Focusing on average effect of lines provided an increase in power to differentiate genotypes.

The differences in transgene effects across hybrids raises a conundrum for selecting candidate transgenes for commercial advancement: how to best to evaluate a transgenes value – and specifically a transgene for a quantitative trait – across the breadth of a breeding program’s germplasm and in a representative set of environments, given that many constructs (and transformation events) will likely need to be evaluated. This study demonstrates that Type II errors can occur even with many reps and locations as a consequence of limited evaluations across genetic backgrounds.

The observed range of transgene effects cautions the likelihood of estimating the value of the transgene based on performance within the context of a single or even a few hybrids, especially if these hybrids always share a single transgenic line as a common parent. In this case, for a transgene that averaged a 0.2 Mg ha^−1^ advantage overall, for nearly all the individual hybrids tested, a non-significant yield difference was observed when compared to the non-transgenic hybrid. Therefore, there would be a high likelihood of discarding the transgene for having no positive effect on yield if only tested in a narrow set of hybrids. This transgene, in specific hybrid combinations, increased yields by up to 8 percent (.8 Mg ha^−1^). By way of comparison, US corn yields have increased by 1 percent per year (USDA-NASS, 2021). Even by considering the general combining ability of transgenic lines based on multiple non-transgenic lines and by testing a non-transgenic line across multiple transgenic lines, we found a similar distribution among and across transgenic lines, which underscores the criticality of selecting an adequate number of transgenic and non-transgenic lines in which to evaluate transgenes affecting highly quantitative traits like yield.

Given a range of transgene effects across germplasm, it is hypothesized that a transgene’s value could be optimized using a forward breeding approach, introducing this gene early in population development and selecting individuals that have strong transgene combining ability. Both phenotypes with significant and non-significant T x G could benefit from such an approach, but for different reasons. For ASI, a phenotype with no significant T x G, the proposition of reducing ASI with a transgene has the potential to expand the breeding germplasm pool that might have otherwise been inappropriate for water-stressed environments. This could increase genetic variance on which to make selections and ultimately increase the rate of genetic gain for yield improvement under such water limited environments. For traits where the transgene effects have large T x G interaction components, selecting the best germplasm to combine with the transgene could significantly improve the chances of successful commercialization. Within the current study lines and hybrid combinations could be selected to maximize the transgene effect for yield while minimizing the effect for plant height.

For the three traits considered in this study there were differences in the interaction components identified. Yield and plant height had significant T x G and T x G x E components, while ASI showed no significant T x G or T x G x E interactions. Further research is required to test the hypothesis that the lack of interactions for ASI is due to the directness of the impact from ethylene on ASI. Overall, the presence of important T x G and T x G x E interactions highlights the risks of Type I and Type II errors and motivates testing of transgenes in larger samples of elite genotypes and relevant environments than is typical in most transgene discovery studies to evaluate their potential efficacy.

## Abbreviations

QTL: Quantitative Trait Locus/Loci

## SUPPLEMENTAL MATERIAL

A supplemental table is included that provides the data from individual hybrid combinations.

## APPENDIX A. STATISTICAL MODELS USED IN ANALYSIS

Model 1 – Used for determining significant interactions:

Y ~ mu Otr Otr.Hetr !f mv !r Fam Fam.Otr Fam.Otr.Hetr Env Env.Otr Env.Otr.Hetr Env.Rep Env.Rep.Fam Env.Fam Env.Fam.Otr Env.Fam.Otr.Hetr at(Env).yadj at(Env).xadj

Y – response variable; mu – overall population mean; Otr – transgene overall with 2 levels (transgenic or non-transgenic); Hetr – transgene effect among heterotic group donor lines with 3 levels (NSS Donor, SS Donor, non-transgenic); Fam – hybrid nest; Env – Environment; Rep – replication; yadj – row level spatial structure; xadj – column level spatial structure

Model 2 – Used for estimating line pairs across non-transgenic lines and non-transgenic across line pairs:

Y ~ mu Otr NSSp SSp NSSp.Otr SSp.Otr !f mv !r Fam Fam.Otr Env Env.Otr Env.Rep Env.Rep.Fam Env.Fam Env.Fam.Otr at(Env).yadj at(Env).xadj

Y – response variable; mu – overall population mean; Otr – transgene overall (transgenic or non-transgenic); NSSp – NSS parent; SSp – SS parent; Fam – hybrid nest; Env – Environment; Rep - replication; yadj – row level spatial structure; xadj – column level spatial structure

