## Supplemental Table 1 for "Transgene by Germplasm Interactions Can Impact Transgene Evaluation"

Supplemental Table 1. Transgene effects for each maize hybrid combination as measured by the difference between the transgenic and non-transgenic hybrid. The transgene donor (NSS or SS) is indicated for each hybrid; a given hybrid combination could have both an SS and a NSS transgene donor. Results are presented over all environments for yield and by stress (FS) and well-watered (WW) environments independently for both traits.

| **Hybrid** | **Yield (Rank)** | | |  | | **Plant Height (cm)** | |
| --- | --- | --- | --- | --- | --- | --- | --- |
| **(Transgene donor)** | **Overall** | **FS^a^** | **WW^b^** | |  | **FS** | **WW** |
|  | ---------------Mg ha^-1^--------------- | | | |  | ---------------cm--------------- | |
| SS1/NSS1 (NSS) ^c^ | 0.0 (65) ^d^ | 0.2 (40) | 0.0 (71) | |  | 4* (11) | 4 (54) |
| SS1/NSS1 (SS) | 0.1 (61) | 0.1 (61) | 0.2 (50) | |  | 2 (62) | 3 (56) |
| SS1/NSS2 (NSS) | 0.0 (71) | 0.0 (62) | 0.1 (63) | |  | 2 (60) | 6 (27) |
| SS1/NSS2 (SS) | 0.0 (66) | -0.1 (73) | 0.3 (36) | |  | 3 (54) | 3 (58) |
| SS1/NSS3 (NSS) | 0.2 (39) | 0.3 (12) | 0.1 (60) | |  | 4* (28) | 6 (36) |
| SS1/NSS3 (SS) | 0.3 (23) | 0.2 (34) | 0.3 (32) | |  | 1 (66) | 6 (38) |
| SS1/NSS4 (NSS) | 0.3 (20) | 0.0 (66) | 0.4 (19) | |  | 4* (17) | 8* (15) |
| SS1/NSS4 (SS) | 0.2 (37) | -0.1 (74) | 0.5* (5) | |  | 0 (75) | 0 (74) |
| SS1/NSS5 (NSS) | 0.1 (58) | 0.3 (13) | -0.1 (74) | |  | 1 (68) | 5 (45) |
| SS1/NSS5 (SS) | 0.0 (68) | 0.2 (36) | 0.1 (67) | |  | 1 (67) | 3 (62) |
| SS1/NSS6 (NSS) | 0.2 (46) | 0.2 (29) | 0.2 (53) | |  | 3 (31) | 6 (25) |
| SS1/NSS6 (SS) | 0.2 (38) | 0.1 (55) | 0.3 (27) | |  | 1 (64) | 1 (72) |
| SS1/NSS7 (SS) | 0.1 (64) | 0.0 (65) | 0.2 (42) | |  | 1 (70) | 1 (71) |
| SS2/NSS1 (NSS) | 0.1 (54) | 0.1 (45) | 0.2 (58) | |  | 4* (16) | 4 (52) |
| SS2/NSS1 (SS) | 0.1 (53) | 0.0 (64) | 0.3 (31) | |  | 3 (51) | 2 (67) |
| SS2/NSS2 (NSS) | 0.1 (55) | 0.3 (17) | 0.1 (68) | |  | 6* (1) | 9* (7) |
| SS2/NSS2 (SS) | 0.2 (35) | 0.1 (43) | 0.2 (44) | |  | 4* (27) | 6* (30) |
| SS2/NSS3 (NSS) | 0.3 (18) | 0.4 (6) | 0.3 (34) | |  | 3* (46) | 8* (10) |
| SS2/NSS3 (SS) | 0.4 (9) | 0.3 (21) | 0.4 (13) | |  | 4* (23) | 2 (68) |
| SS2/NSS4 (NSS) | 0.3 (21) | -0.1 (72) | 0.5 (10) | |  | 4* (25) | 3 (61) |
| SS2/NSS4 (SS) | 0.1 (60) | -0.2 (77) | 0.6* (1) | |  | 2 (63) | 2 (66) |
| SS2/NSS5 (NSS) | -0.4 (77) | 0.1 (51) | -0.4 (77) | |  | 1 (72) | 2 (65) |
| SS2/NSS5 (SS) | -0.2 (76) | 0.0 (69) | -0.2 (76) | |  | 4* (18) | 5* (43) |
| SS2/NSS6 (NSS) | 0.1 (57) | 0.3 (11) | 0.1 (69) | |  | 4* (24) | 7* (20) |
| SS2/NSS6 (SS) | 0.3 (27) | 0.2 (31) | 0.2 (47) | |  | 3 (32) | 5* (40) |
| SS2/NSS7 (SS) | 0.3 (26) | 0.1 (60) | 0.4 (12) | |  | 3 (58) | 1 (70) |
| SS3/NSS1 (NSS) | 0.2 (44) | 0.2 (27) | 0.2 (57) | |  | 3 (40) | 8* (11) |
| SS3/NSS2 (NSS) | 0.2 (47) | 0.2 (38) | 0.2 (49) | |  | 4* (10) | 7* (19) |
| SS3/NSS3 (NSS) | 0.2 (43) | 0.3 (18) | 0.1 (59) | |  | 5* (3) | 10* (2) |
| SS3/NSS4 (NSS) | 0.7* (2) | 0.8* (1) | 0.3 (26) | |  | 4* (12) | 10* (5) |
| SS3/NSS5 (NSS) | -0.2 (75) | 0.1 (54) | -0.1 (75) | |  | 0 (73) | 4 (53) |
| SS3/NSS6 (NSS) | 0.4 (10) | 0.5 (4) | 0.2 (43) | |  | 5* (2) | 11* (1) |
| SS4/NSS1 (NSS) | 0.2 (45) | 0.3 (19) | 0.1 (61) | |  | 5* (7) | 7* (16) |
| SS4/NSS2 (NSS) | 0.8* (1) | 0.7* (2) | 0.4 (18) | |  | 5* (6) | 10* (3) |
| SS4/NSS3 (NSS) | 0.3 (28) | 0.2 (28) | 0.3 (33) | |  | 4* (15) | 6* (31) |
| SS4/NSS4 (NSS) | 0.7* (3) | 0.4 (8) | 0.6* (3) | |  | 2 (59) | 8* (13) |
| SS4/NSS5 (NSS) | 0.0 (69) | 0.0 (70) | 0.2 (52) | |  | 2 (61) | 3 (60) |
| SS4/NSS6 (NSS) | 0.1 (59) | 0.1 (49) | 0.2 (51) | |  | 4* (26) | 7* (17) |
| SS5/NSS1 (NSS) | 0.0 (72) | 0.3 (23) | 0.0 (72) | |  | 3 (41) | 10* (4) |
| SS5/NSS1 (SS) | 0.1 (50) | 0.1 (48) | 0.1 (62) | |  | 3 (42) | 8* (12) |
| SS5/NSS2 (NSS) | 0.2 (40) | 0.3 (14) | 0.2 (55) | |  | 3 (39) | 3 (55) |
| SS5/NSS2 (SS) | 0.3 (22) | 0.2 (39) | 0.3 (29) | |  | 3 (45) | 5 (41) |
| SS5/NSS3 (NSS) | 0.2 (33) | 0.2 (32) | 0.2 (45) | |  | 4* (21) | 6 (32) |
| SS5/NSS3 (SS) | 0.2 (36) | 0.1 (56) | 0.4 (20) | |  | 4* (22) | 6 (34) |
| SS5/NSS4 (NSS) | 0.4 (11) | 0.2 (33) | 0.4 (11) | |  | 3 (44) | 6 (26) |
| SS5/NSS4 (SS) | 0.4 (12) | 0.1 (57) | 0.6* (2) | |  | 3 (49) | 5 (42) |
| SS5/NSS5 (NSS) | 0.2 (41) | 0.3 (20) | 0.2 (56) | |  | -3 (77) | - |
| SS5/NSS5 (SS) | 0.3 (30) | 0.1 (46) | 0.3 (30) | |  | 3 (57) | 4 (46) |
| SS5/NSS6 (NSS) | 0.0 (67) | 0.1 (44) | 0.1 (66) | |  | 3 (48) | 5 (39) |
| SS5/NSS6 (SS) | 0.1 (51) | 0.0 (63) | 0.2 (40) | |  | 4* (8) | 6 (29) |
| SS5/NSS7 (SS) | 0.4 (13) | 0.3 (15) | 0.4 (22) | |  | 3* (29) | 4 (49) |
| SS6/NSS1 (NSS) | 0.1 (63) | 0.0 (67) | 0.2 (38) | |  | 3 (56) | 7* (21) |
| SS6/NSS1 (SS) | 0.1 (48) | -0.1 (75) | 0.4 (15) | |  | 1 (65) | 6* (35) |
| SS6/NSS2 (NSS) | 0.2 (34) | 0 (71) | 0.3 (25) | |  | 3 (30) | 9* (8) |
| SS6/NSS2 (SS) | 0.1 (56) | -0.2 (76) | 0.5 (9) | |  | 3 (50) | 6 (28) |
| SS6/NSS3 (NSS) | 0.1 (52) | 0.3 (22) | 0.1 (70) | |  | 4* (13) | 7 (22) |
| SS6/NSS3 (SS) | 0.1 (49) | 0.1 (47) | 0.2 (48) | |  | 3 (35) | 4 (48) |
| SS6/NSS4 (NSS) | 0.5 (6) | 0.2 (26) | 0.4 (16) | |  | 3 (34) | 7* (18) |
| SS6/NSS4 (SS) | 0.4 (14) | 0.1 (53) | 0.6* (4) | |  | 0 (74) | 4 (47) |
| SS6/NSS5 (NSS) | -0.1 (74) | 0.1 (50) | -0.1 (73) | |  | 1 (69) | 0 (73) |
| SS6/NSS5 (SS) | -0.1 (73) | 0.0 (68) | 0.1 (65) | |  | 1 (71) | -1 (76) |
| SS6/NSS6 (NSS) | 0.1 (62) | 0.3 (16) | 0.1 (64) | |  | 3 (36) | 9* (6) |
| SS6/NSS6 (SS) | 0.3 (19) | 0.2 (42) | 0.3 (37) | |  | 3 (55) | 6 (37) |
| SS6/NSS7 (SS) | 0.2 (42) | 0.2 (41) | 0.2 (39) | |  | 3 (47) | 7* (24) |
| SS7/NSS1 (NSS) | 0.5 (8) | 0.6 (3) | 0.3 (24) | |  | 5* (5) | 5 (44) |
| SS7/NSS1 (SS) | 0.7* (4) | 0.5 (5) | 0.5 (8) | |  | 3 (43) | -1 (75) |
| SS7/NSS2 (NSS) | 0.3 (25) | 0.2 (25) | 0.3 (35) | |  | 4* (9) | 8* (9) |
| SS7/NSS2 (SS) | 0.3 (29) | 0.1 (52) | 0.4 (14) | |  | 3 (38) | 7* (23) |
| SS7/NSS3 (NSS) | 0.2 (31) | 0.2 (35) | 0.2 (46) | |  | 3 (37) | 8* (14) |
| SS7/NSS3 (SS) | 0.2 (32) | 0.1 (58) | 0.4 (21) | |  | 3 (52) | 3 (57) |
| SS7/NSS4 (NSS) | 0.0 (70) | 0.2 (37) | 0.2 (54) | |  | 3 (53) | 3 (59) |
| SS7/NSS4 (SS) | 0.3 (17) | 0.1 (59) | 0.3 (28) | |  | 4* (19) | 2 (64) |
| SS7/NSS5 (NSS) | 0.3 (24) | 0.3 (10) | 0.4 (23) | |  | 0 (76) | 2 (69) |
| SS7/NSS5 (SS) | 0.5* (7) | 0.2 (30) | 0.5* (7) | |  | 3* (33) | 4 (51) |
| SS7/NSS6 (NSS) | 0.3 (16) | 0.4 (7) | 0.2 (41) | |  | 5* (4) | 6* (33) |
| SS7/NSS6 (SS) | 0.4 (15) | 0.2 (24) | 0.4 (17) | |  | 4* (14) | 3 (63) |
| SS7/NSS7 (SS) | 0.6* (5) | 0.3 (9) | 0.5 (6) | |  | 4* (20) | 4 (50) |

^a^FS=Flowering stress, ^b^WW=Well-watered, ^c^NSS=Non Stiff Stalk, SS=Stiff Stalk, ^d^Difference between non-transgenic and transgenic hybrids for a given non-transgenic line is significant (*) at the 5% probability level.
